# Transcriptional signatures of schizophrenia in hiPSC-derived NPCs and neurons are concordant with signatures from post mortem adult brains

**DOI:** 10.1101/185546

**Authors:** Gabriel E. Hoffman, Brigham J. Hartley, Erin Flaherty, Ian Ladran, Peter Gochman, Doug Ruderfer, Eli A. Stahl, Judith Rapoport, Pamela Sklar, Kristen J. Brennand

## Abstract

Whereas highly penetrant variants have proven well-suited to human induced pluripotent stem cell (hiPSC)-based models, the power of hiPSC-based studies to resolve the much smaller effects of common variants within the size of cohorts that can be realistically assembled remains uncertain. In developing a large case/control schizophrenia (SZ) hiPSC-derived cohort of neural progenitor cells and neurons, we identified and accounted for a variety of technical and biological sources of variation. Reducing the stochastic effects of the differentiation process by correcting for cell type composition boosted the SZ signal in hiPSC-based models and increased the concordance with post mortem datasets. Because this concordance was strongest in hiPSC-neurons, it suggests that this cell type may better model genetic risk for SZ. We predict a growing convergence between hiPSC and post mortem studies as both approaches expand to larger cohort sizes. For studies of complex genetic disorders, to maximize the power of hiPSC cohorts currently feasible, in most cases and whenever possible, we recommend expanding the number of individuals even at the expense of the number of replicate hiPSC clones.

## INTRODUCTION

A growing number of studies have demonstrated that human induced pluripotent stem cells (hiPSCs) can serve as cellular models of both syndromic ^1-5^ and idiopathic ^6-11^ forms of a variety of neurodevelopmental disorders. We and others have previously shown that hiPSC-derived neural progenitor cells (NPCs) and neurons generated from patients with schizophrenia (SZ) show altered gene and microRNA expression ^4,10-15^, which may underlie observed *in vitro* phenotypes such as aberrant hiPSC-NPC polarity ^5^ and migration ^13,16^, as well as deficits in hiPSC-neuron connectivity and function ^11,17-19^. Altogether, such hiPSC-based approaches seem to capture aspects of SZ biology identified through post mortem studies and animal models ^20^. Nonetheless, mechanistic studies to date have tended to focus on rare variants ^4,5,19^; the ability of an hiPSC-based approach to resolve the much smaller effects of common variants remained uncertain.

We established a case-control SZ cohort structure designed to capture a broad range of rare and common variants that might underlie SZ risk, in order to address and quantify the intra-and inter-individual variability inherent in this approach and uncover to what extent hiPSC-based models can identify common pathways underlying such different genetic risk factors (**Fig. 1**). Because hiPSC-neurons are likely best suited for the study of disease predisposition ^13,21-24^, we applied this methodology to a childhood-onset SZ (COS) cohort, a subset of SZ patients defined by onset, severity and prognosis ^25-27^. COS patients have a more salient genetic risk, with a higher rate of SZ-associated copy number variants (CNVs) ^28^ and stronger common SZ polygenic risk scores ^29^. Overall, across 94 RNA-Seq samples, we observed many sources of variation reflecting both biological (i.e. reprogramming and differentiation) and technical effects. By systematically accounting for covariates and adjusting for heterogeneity in neural differentiation, we improved our ability to resolve the disease-relevant signal. Our bioinformatic pipeline reduces the risk of false positives arising from the small sample sizes of hiPSC-based approaches and we hope it can help guide data analysis in similar hiPSC-based disease studies.

**Figure 1:**
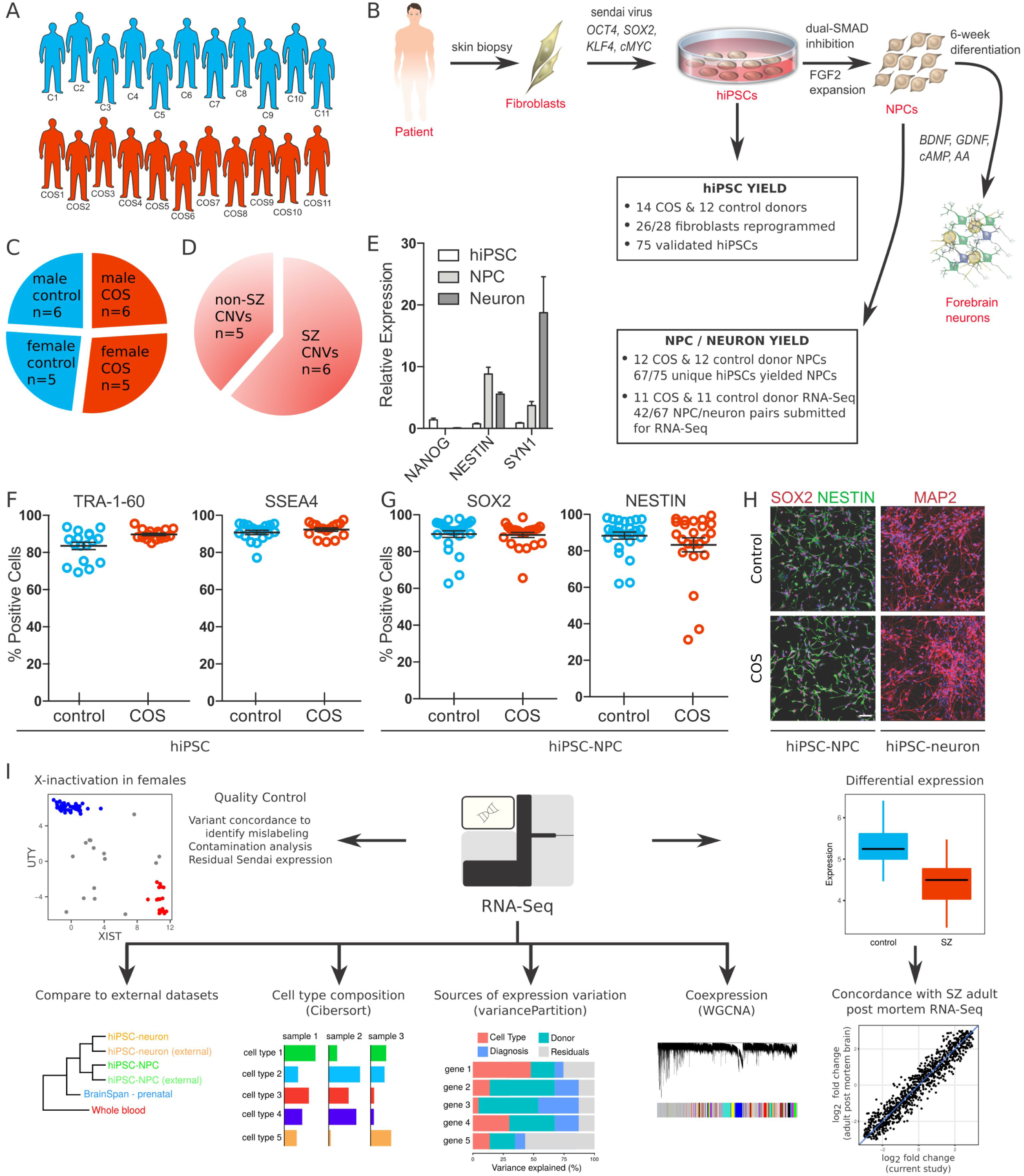
COS hiPSC cohort reprogramming and differentiation. **A)** Validated hiPSCs (from 14 individuals with childhood-onset-schizophrenia (COS) and 12 unrelated healthy controls) and NPCs (12 COS; 12 control individuals) yielded 94 RNA-Seq samples (11 COS; 11 control individuals). **B)** Schematic illustration of the reprogramming and differentiation process, noting the yield at each stage. **C)** Sex breakdown of the COS-control cohort. **D)** Breakdown of SZ-associated copy number variants in the 11 COS patients with RNA-Seq data. **E)** Representative qPCR validation of *NANOG, NESTIN* and *SYN1* expression in hiPSCs (white bar), NPCs (light grey) and 6-week-old neurons (dark grey) from three individuals. **F**) FACS analysis for pluripotency markers TRA-1-60 (left) and SSEA4 (right) in representative control (blue, n=17) and COS (red, n=16) hiPSCs. **G**) FACS analysis for NPC markers SOX2 (left) and NESTIN (right) in control (blue, n=34) and COS (red, n=37) NPCs. **H**) Representative images of NPCs (left) and 6-week-old forebrain neurons (right) from control (top) and COS (bottom). NPCs stained with SOX2 (red) and NESTIN (green); neurons stained with MAP2 (red). DAPI-stained nuclei (blue). Scale bar 50 ?m. **I**) Computational workflow showing quality control, integration with external datasets, computational deconvolution with Cibersort, decomposition multiple sources of expression variation with variancePartition, coexpression analysis with WGCNA, differential expression and concordance analysis.

## RESULTS

### Generation, validation and transcriptomic profiling of a large cohort of COS hiPSC-NPCs and hiPSC-neurons

Individuals with COS as well as unaffected, unrelated healthy controls were recruited as part of a longitudinal study conducted at the National Institute of Health ^28,29^ (see **SI Table 1** for available clinical information). This cohort is comprised of nearly equal numbers of cases and controls (**Fig. 1A,B,C**); 16 cases were selected representing a range of SZ-relevant CNVs, including 22q11.2 deletion, 16p11.2 duplication, 15q11.2 deletion and *NRXN1* deletion (2p16.3) ^30^ and/or idiopathic genetics with a strong family history of SZ, 12 controls were identified as being most appropriately matched for sex, age and ethnicity (**Fig. 1D;** **SI Table 1**).

We used an integration free approach to generate genetically unmanipulated hiPSCs from COS patients (14 of 16 patients, 88% reprogrammed) and unrelated age-and sex-matched controls (12 of 12 controls, 100% reprogrammed) (**Fig. 1B**). Briefly, primary fibroblasts were reprogrammed by sendai viral delivery of *KLF4, OCT4, SOX2* and *cMYC;* presumably clonal lines were picked and expanded 23-30 days following transduction. Following extensive immunohistochemistry, florescent activated cell sorting (FACS), quantitative polymerase chain reaction (qPCR) and karyotype assays to assess the quality of the hiPSCs (**Fig. 1B,E,F**), we selected two to three presumably clonal hiPSC lines per individual (n=40 COS, n=35 control, **Table 1; SI Table 1**). A subset of these hiPSCs has been previously reported ^10,31^.

**Table 1.** Number of individuals and cell lines at each step of experimental workflow.

| Experimental workflow | Total individuals |  | Total hiPSC lines |  | Total NPC lines |  | Total Neurons |  |
| --- | --- | --- | --- | --- | --- | --- | --- | --- |
|  | control | COS | control | COS | control | COS | control | COS |
| Fibroblasts | 12 | 16 | - | - | - | - | - | - |
| hiPSCs | 12 | 14 | 35 | 40 | - | - | - | - |
| NPCs | 12 | 12 | 35 | 32 | 35 | 32 | - | - |
| RNA submitted | 11 | 11 | 20 | 22 | 24 | 23 | 24 | 23 |
| RNA-Seq QC passed | 9 | 10 | 17 | 18 | 20 | 18 | 20 | 18 |

Using dual-SMAD inhibition ^32^, three to five forebrain hiPSC-NPC populations were differentiated from each validated hiPSC line via an embryoid body intermediate, once hiPSCs had been passaged approximately ten times. hiPSC-hiPSC-NPCs with normal morphology and robust protein levels of NESTIN and SOX2 by FACS and/or immunocytochemistry (**Fig. 1G,H**) (n=32 COS, n=35 control hiPSC-NPCs representing 67 unique hiPSC lines reprogrammed from 12 unique COS and 12 unique control individuals) were selected for further differentiation to 6-week-old forebrain neuronal populations (**Table 1**; **SI Table 2**). We ^11,13,33^ and others ^1,19,34^ have previously demonstrated that hiPSC-NPCs can be directed to differentiate into mixed populations of excitatory neurons, inhibitory neurons and astrocytes. hiPSC-neurons have neuronal morphology, undergo action potentials, release neurotransmitters, show evidence of spontaneous synaptic activity, and resemble the gene expression of fetal forebrain tissue.

Because it required nearly four years to generate and differentiate all hiPSCs, hiPSC-NPCs and hiPSC-neurons, it was not possible to fully apply standardized conditions across all cellular reprogramming and neural differentiations. Media reagents, substrates and growth factors for fibroblast expansion, reprogramming, hiPSC differentiation, NPC expansion and neuronal differentiation, as well as personnel and laboratory spaces, varied over time. While individual fibroblast lines were reprogrammed and differentiated to hiPSC-NPCs in the order in which they were received, multiple randomization steps were introduced at the subsequent stages, particularly the thaw, expansion, and neuronal differentiation of validated hiPSC-NPCs in preparation for RNA sequencing (RNA-Seq) (see **SI Table 2** for available batch information). Only validated hiPSC-NPCs that yielded high quality populations of matched hiPSC-NPCs and hiPSC-neurons in one of three batches of thaws were used for RNA-Seq (**SI Table 1,2**).

RNA-Seq data was generated from 94 samples (n=47 hiPSC-NPC, n=47 hiPSC-neurons; n=46 COS, n=48 controls; representing 42 unique hiPSC lines reprogrammed from 11 unique COS and 11 unique control individuals) following ribosomal RNA (rRNA) depletion (**Table 1**; **SI Table 2**). The median number of uniquely mapped read pairs per sample was 42.7 million, of which only a very small fraction were rRNA reads (**SI Fig. 1; SI Table 3**). 18,910 genes (based on ENSEMBL v70 annotations) were expressed at levels deemed sufficient for analysis (at least 1 CPM in at least 30% of samples); 11,681 were protein coding, 879 were lincRNA, and the remaining were of various biotypes (**SI Table 4**).

Since six COS patients were selected based on CNV status, we examined gene expression in the regions affected by the CNVs. Despite the noise inherent to RNA-Seq and the high level of biologically driven expression variation in samples without CNVs, we identified corresponding hiPSC-NPC and neuron expression changes in some CNV regions (**SI Fig. 2**).

In addition to SZ diagnosis-dependent effects, gene expression between hiPSC-NPCs and hiPSC-neurons was expected to vary as a result of technical ^35^, epigenetic ^36-38^ and genetic ^39-41^ differences ^42^. Unexpectedly, we also observed substantial variation in cell type composition (CTC) between populations of hiPSC-NPCs and hiPSC-neurons. In the following sections, we discuss our strategy to address these sources of variation (**Figs. 2**-**4**).

**Figure 2:**
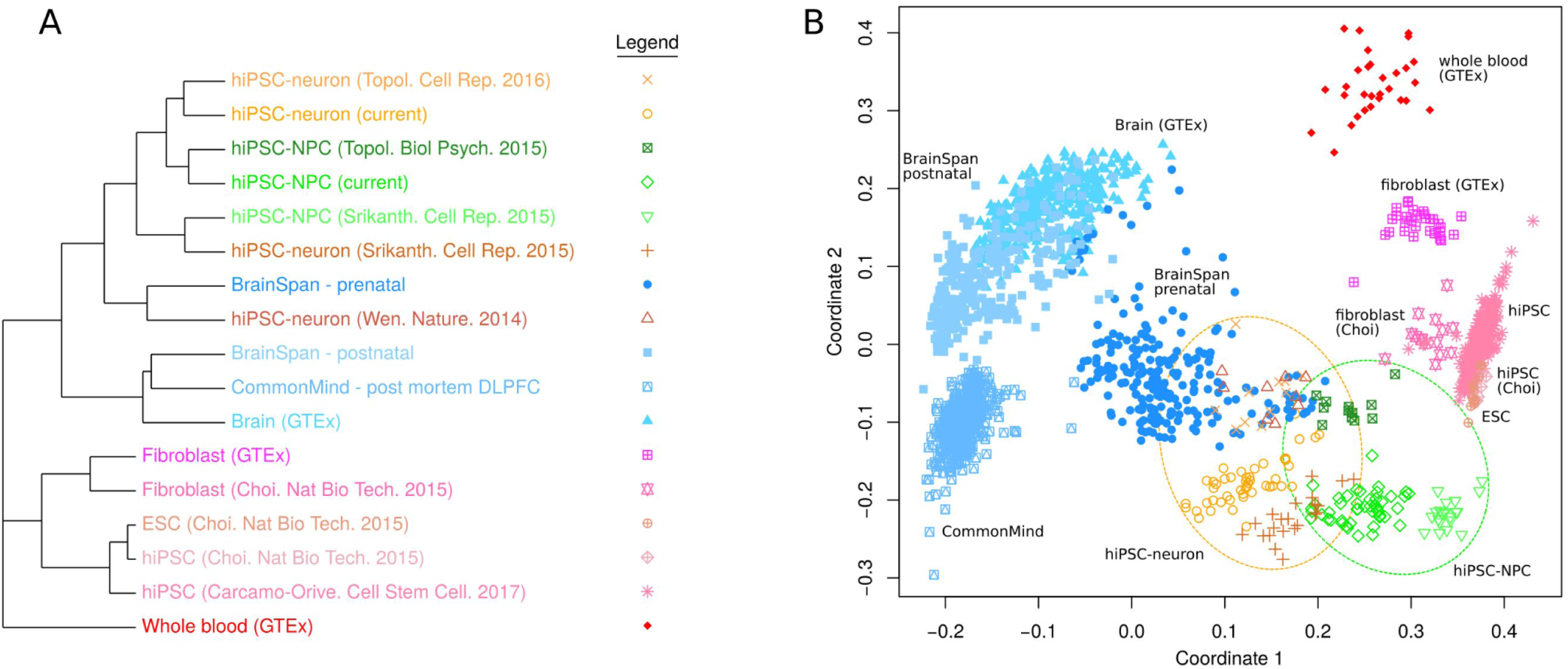
Cell type specificity of gene expression. **A**) Summary of hierarchical clustering of 2082 RNA-Seq samples shows clustering by cell type. A pairwise distance matrix was computed for all samples, and the median distance between all samples in each category were used to create a summary distance matrix using to perform the final clustering. **B**) Multidimensional scaling with samples colored as in (A). hiPSC-NPCs from multiple studies are indicated in the green circle, and hiPSC-neurons from multiple studies are indicated in the orange circle.

**Figure 3:**
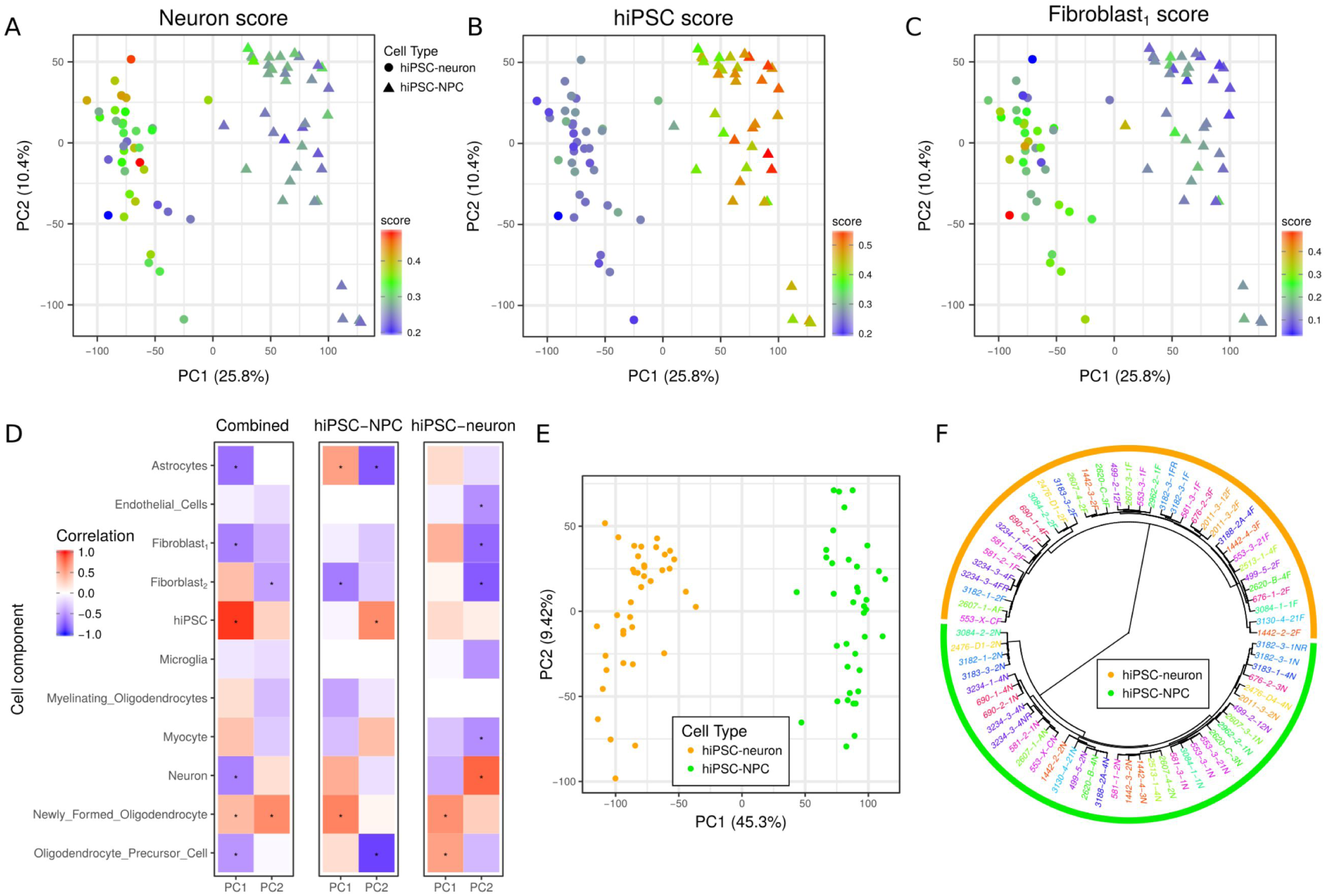
Variation in cell type composition contributes to gene expression variation. **A,B,C**) Principal components analysis of gene expression data from hiPSC-NPCs (triangles) and hiPSC-neurons (circles) where samples are colored according to their cell type composition scores from cibersort for A) neuron, B) hiPSC, and C) fibroblast_1_ components. Color gradient is shown on the bottom right of each panel. **D**) Correlation between 11 cell type composition scores for the first two principal components of gene expression data from all samples, only hiPSC-NPCs, and only hiPSC-neurons. Red indicates a strong positive correlation with a principal component and blue indicates a strong negative correlation. Asterisks indicate correlations that are significantly different from zero with a p-value that passes the Bonferroni cutoff of 5% for 66 tests. **E**) Principal components analysis of expression residuals after correcting for the two fibroblast cell type composition scores. **F**) Hierarchical clustering of samples based on expression residuals after correcting for the two fibroblast cell type composition scores.

**Figure 4:**
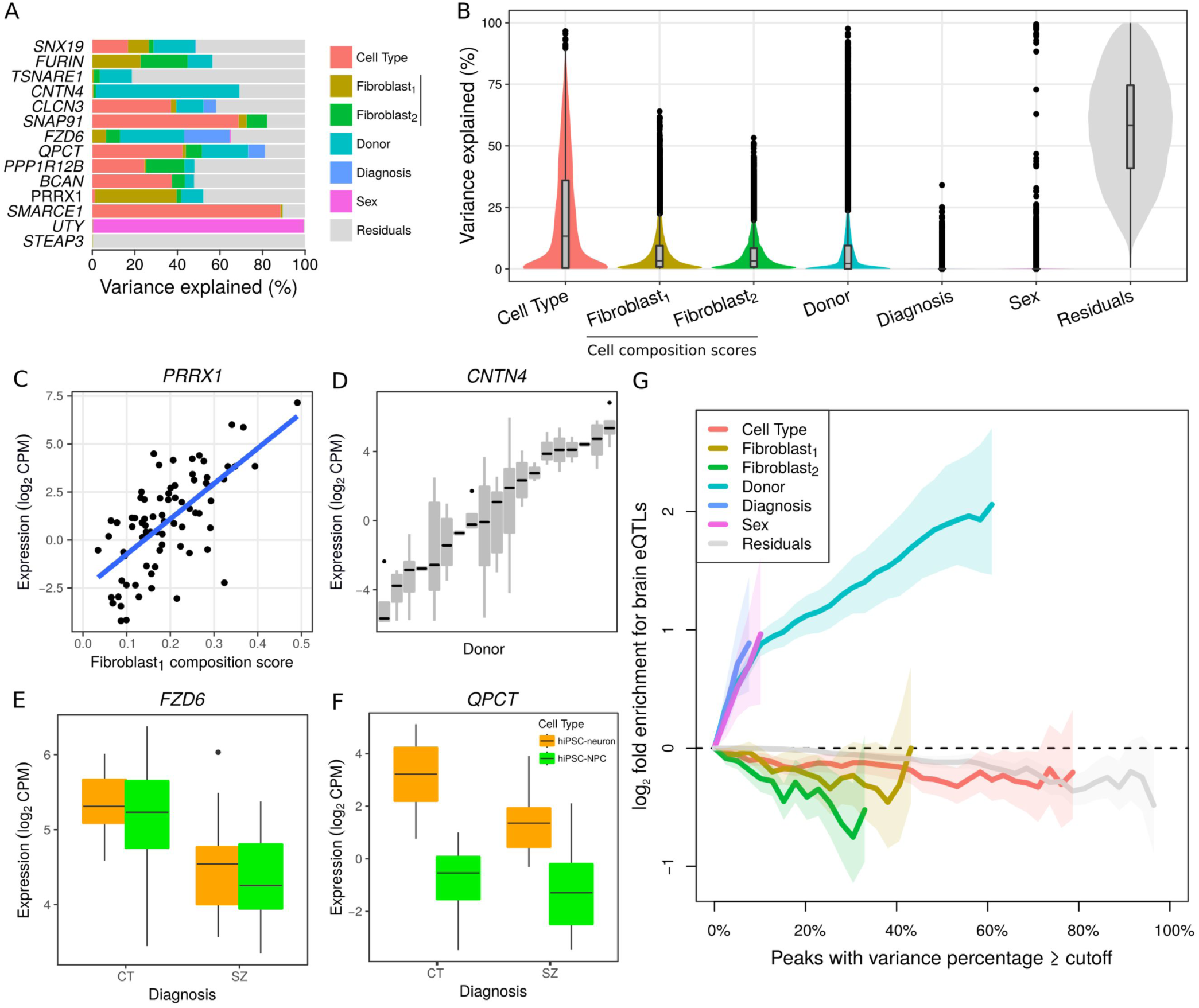
Decomposing expression variation into multiple sources. **A**) Expression variance is partitioned into fractions attributable to each experimental variable. Genes shown include genes of known biological relevance to schizophrenia and genes for which one of the variables explains a large fraction of total variance. **B**) Violin plots of the percentage of variance explained by each variable over all the genes. **C**-**F**) Expression of representative genes stratified by a variable that explains a substantial fraction of the expression variation. C) *PRRX1* plotted as a function of the fibroblast_1_ cell type composition score. D) *CNTN4* stratified by Donor. E) *FDZ6* stratified by disease status and cell type. F) *QPCT* stratified by disease status and cell type. **G**) Genes that vary most across donors are enriched for brain cis-eQTLs. Fold enrichment (log_2_) for the 2000 top cis-eQTLs discovered in post mortem dorsolateral prefrontal cortex data generated by the CommonMind Consortium^48^ shown for six sources of variation, plus residuals. Each line indicates the fold enrichment for genes with the fraction of variance explained exceeding the cutoff indicated on the x-axis. Shaded regions indicate the 90% confidence interval based on 10,000 permutations of the variance fractions. Enrichments are shown on the x-axis until less that 100 genes pass the cutoff.

### Addressing technical variation in hiPSC-NPC and neuron RNA-Seq data

We implemented an extensive quality control pipeline to detect, minimize and account for many possible sources of technical variation (**Fig. 1I**). Samples were submitted and processed for RNA-Seq in only one batch; RNA isolation, library preparation and sequencing were completed under standardized conditions at the New York Genome Center. Errors in sample mislabeling and cell culture contamination were identified, allowing us to correct sample labeling when possible and remove samples from further analysis when not. Batch effects in both tissue culture and RNA-Seq sample processing were corrected for and samples with aberrant X-inactivation ^43^ and/or residual Sendai virus expression were flagged.

Expression patterns of genes on the sex chromosomes can identify the sex of each sample, confirm sample identity, and also measure the extent of X-inactivation in females. Using *XIST* on chrX and the expression of six genes on chrY (*USP9Y*, *UTY*, *NLGN4Y*, *ZFY*, *RPS4Y1*, *TXLNG2P*), this analysis identified 2 mislabeled males that show a female expression pattern and 15 female samples that have expression patterns intermediate between males and females (**SI Fig. 3A**), consistent with either contamination or aberrant X-inactivation.

Samples with mislabeling and/or cross-individual contamination, whether during cell culture and/or RNA library preparation, were identified through genotype concordance analysis. VerifyBamID ^44^ was used to compare the genotype of the parental fibroblast samples with variants called from RNA-Seq data from the respective hiPSC-NPCs and hiPSC-neurons. In total, 76 samples (81%; n=38 hiPSC-NPC, n=38 hiPSC-neurons; n=36 COS, n=40 controls, from 10 unique COS and 9 unique control individuals) were validated for subsequent analysis (**Table 1**; **SI Table 2**; **SI Fig. 3B**).

Residual Sendai virus expression was assessed using Inchworm in the Trinity package ^45^, which performed *de novo* assembly of reads that did not map to the human genome. Comparisons of these contigs to the Sendai virus genome sequence (GenBank: AB855655.1) quantified the number of reads corresponding to residual Sendai expression in each NPC and neuron sample. Although Sendai viral vectors are widely assumed to be lost within eleven hiPSC passages ^46^, and that on average our hiPSCs were passaged >10-15 times and our hiPSC-NPCs >5 times, we identified Sendai viral transcripts in a subset of our samples. While the majority (70 of 87, 80%) (75 of the total 94, 79.8%) of RNA-Seq samples did not contain any reads that mapped to the Sendai viral genome, 17 (or 19 of total) samples (**SI Table 2; SI Fig. 4**) showed evidence of persistent Sendai viral expression at > 1 count per million. Differential expression analysis identified 2768 genes correlated with Sendai expression at FDR < 5% (**SI Table 5**). We note that this signal is not driven by outliers since quantile normalized Sendai expression values were used in this analysis. In fact, these genes are highly enriched for targets of *MYC* (OR = 3.75, p < 6.4e-38) (**SI Table 6**, **SI Fig. 5A**). Although *MYC* is one of the four transcription factors (along with *SOX2, KLF4*, and *OCT4)* used in hiPSC reprogramming, expression of these four genes was not associated with Sendai expression (**SI Fig. 5B)**. The correlation of residual Sendai expression with activation of *MYC* targets suggests that this could be a potential source of transcriptional and phenotypic variation in hiPSCs; however, neither incorporating Sendai expression as a covariate nor dropping samples with Sendai expression from downstream expression meaningfully impacted overall findings.

Overall, our rigorous bioinformatic strategy adjusted for technical variation and batch effects, eliminated spurious samples, and flagged samples that were contaminated or had aberrant X-inactivation. This extensive analysis was motivated by the high level of intra-donor expression variation (see below), and eliminating these factors as possible explanations for this expression variation ultimately improved our ability to resolve SZ-relevant biology in our dataset.

### COS hiPSC-NPC and hiPSC-neuron RNA-Seq data cluster with existing hiPSC and post mortem brain datasets

To assess the similarity of our hiPSC-NPCs and hiPSC-neurons to other hiPSC studies (by ourselves and others), as well as to post mortem brain, we compared our dataset to publically available hiPSC, hiPSC-derived NPCs/neurons, and post mortem brain homogenate expression data sets (**Fig. 2**). Hierarchical clustering indicated that similarity in expression profiles is largely determined by cell type (**Fig. 2A**). hiPSC-NPC and hiPSC-neuron datasets were more similar to prenatal samples than postnatal or adult post mortem samples ^47-49^, which is consistent with previous reports ^13,21-24^. hiPSC-NPCs and hiPSC-neurons, as well as post mortem brain samples, cluster separately from hiPSCs, ESCs, fibroblasts and whole blood ^35,47,50^. Despite being reprogrammed and differentiated through different methodologies, hiPSC-NPCs and hiPSC-neurons from the current study cluster with hiPSC-NPCs and hiPSC-neurons, respectively, generated previously in the same lab ^10,12^ and with hiPSC-NPCs and hiPSC-neurons from others ^14^, although some hiPSC-neurons ^19^ are more similar to prenatal brain samples from multiple brain regions ^49^. Consistent with a differentiation paradigm from hiPSC to NPC to neuron, multidimensional scaling analysis (**Fig. 2B**) indicated that hiPSC-NPCs more resemble hiPSCs / hESCs than do hiPSC-neurons.

Genome-wide, hiPSC-NPCs and hiPSC-neurons express a common set of genes, so that expression differences between these cell types are driven by changes in expression magnitude rather than activation of entirely different transcriptional modules (**SI Fig. 6**). Moreover, for both hiPSC-NPCs and hiPSC-neurons, genes that show high variance across donors in each cell type are enriched for brain eQTLs (**SI Fig. 7**). Taken together, these two insights justified case-control comparisons within and between both hiPSC-NPCs and hiPSC-neurons.

### Large heterogeneity in cell type composition in both COS and control hiPSC-NPCs and hiPSC-neurons

Given the substantial variability we observed between hiPSC-NPCs and hiPSC-neurons, even from the same individual (**SI Fig. 8**), it seemed likely that inter-hiPSC and inter-NPC differences in differentiation propensity led to unique neural compositions in each sample. hiPSC-NPCs show extensive cell-to-cell variation in the expression of forebrain and neural stem cell markers ^13^ and 6-week-old neurons are comprised of a heterogeneous mixture of predominantly excitatory neurons, but also inhibitory and rare dopaminergic neurons, as well as astrocytes ^11^. We hypothesized that CTC could be inferred using existing single cell RNA-Seq datasets and would enable us to (partially) correct for variation in differentiation efficiencies and account for some of the intra-individual expression variation.

Bulk RNA-Seq analysis reflects multiple constituent cell types; therefore, we performed computational deconvolution analysis using CIBERSORT ^51^ to estimate CTC scores for each hiPSC-NPC and hiPSC-neuron sample (**Fig. 3**). A reference panel of single cell sequencing data from mouse brain ^52^, mouse cell culture of single neural cells ^53^ and bulk RNA-Seq from hiPSC ^50^ was applied.

Overlaying CTC scores on a principal component analysis (PCA) of the expression data indicates that hiPSC-NPCs and hiPSC-neurons separate along the first principal component (PC), explaining 37.6% of the variance, and that the cell types have distinct CTC scores (**Fig. 3A-C**). As expected, hiPSC-neuron samples had a higher neuron CTC score than hiPSC-NPCs (**Fig. 3A**), while hiPSC-NPCs had a higher hiPSC CTC score, consistent with a “stemness” signal (a neural stem cell profile was lacking from our reference) (**Fig. 3B**). Unexpectedly, hiPSC-NPCs had a higher fibroblast_1_ score (**Fig. 3C**). Rather than imply that there are functional fibroblasts within the hiPSC-NPC populations, we instead posit that this fibroblast signature is instead marking a subset of unpatterned, potentially non-neuronal cells ^53^. Analysis of external NPC and neuron datasets indicates that these observations were reproducible, although there is substantial variability in CTC scores across datasets (**SI Fig. 9**). Although correction for CTC improved our ability to distinguish hiPSC-NPC and hiPSC-neuron populations, nonetheless, there remained substantial variability within both the hiPSC-NPCs and hiPSC-neurons that corresponded to CTC (**Fig. 3D**).

The effect of CTC heterogeneity, likely due to the variation in differentiation efficiency, can be reduced by including multiple CTC scores in a regression model and computing the residuals. Using an unbiased strategy, we systematically evaluated which CTC score(s), when included in our model, most explained the variance in our samples. PCA on the residuals from a model including fibroblast_1_ and fibroblast_2_ CTC scores showed a markedly greater distinction between cell types, such that the first PC now explained 45.3% of the variance (**Fig. 3E**). Moreover, accounting for the CTC scores increased the similarity between the multiple biological replicates generated from the same donor and resulted in less intra-individual variation within each cell type (**Fig. 3F**, **SI Fig. 10**). Finally, accounting for CTC was necessary in order to see concordance with one of the adult post mortem cohorts (see below).

### Characterizing known sources of expression variation in COS and control NPC and neuron RNA-Seq dataset

As discussed above, gene expression (in our dataset and others) is impacted by a number of biological and technical factors. By properly attributing multiple sources of expression variation, it is possible to (partially) correct for some variables. To decompose gene expression into the percentage attributable to multiple biological and technical sources of variation, we applied variancePartition ^54^ (**Fig. 4**). For each gene we calculated the percentage of expression variation attributable to cell type, donor, diagnosis, sex, as well as CTC scores for both fibroblast sets. All remaining expression variation not attributable to these factors was termed residual variation. The influence of each factor varies widely across genes; while expression variation in some genes is attributable to cell type, other genes are affected by multiple factors (**Fig. 4A**). Overall, and consistent with the separation of hiPSC-NPCs and hiPSC-neurons by the first PC, cell type has the largest genome-wide effect and explained a median of 13.3% of the observed expression variation (**Fig. 4B**). Expression variation due to diagnosis (i.e. between SZ and controls) had a detectable effect in a small number of genes. Meanwhile, variation across the sexes was small genome-wide, but it explained a large percentage of expression variation for genes on chrX and chrY. Technical variables such as hiPSC technician, hiPSC date, NPC generation batch, NPC technician, sample name, NPC thaw and RIN explained little expression variation (**SI Fig. 11**), especially compared to technical effects observed in previous studies ^35,54^.

Variation attributable to cell type heterogeneity across the CTC scores had a larger median effect than the variation across the 22 donors (fibroblast_1_: 3.3%, fibroblast_2_: 3.2%). The median observed variation across donor is 2.2%, substantially lower than reported in other datasets from hiPSCs ^35,55^ and other cell types ^54^. By considering CTC in our model, the percentage of variation explained by donor significantly increased (median increase to 2.4%, p < 5.8e-62, one-sided paired Wilcoxon), indicating that cell type heterogeneity is an important source of intra-donor expression variation that obscures some inter-donor variation (i.e. case/control differences) of particular biological interest. Critically, there is no apparent diagnosis dependent variation in CTC (**SI Fig. 12**). By compensating for CTC, we prevent variation in neuronal differentiation between hiPSCs from overriding some of the donor-specific gene expression signature that is the central focus of patient-derived cell culture models.

The percentage of expression variation explained by each factor has a specific biological interpretation. *PRRX1* is known to function in fibroblasts ^56,57^ and variation in the fibroblast_1_ CTC score explains 38.3% of expression variant in this gene (**Fig. 4C**). Expression of *CNTC4* is driven by an eQTL in brain tissue that corresponds a risk locus for schizophrenia ^48^. In our data, *CNTC4* has 67.4% expression variation across donors suggesting that this variation is driven by genetics (**Fig. 4D**). Genes that vary across diagnosis correspond to differentially expressed genes, including *FZD6*, a WNT signaling gene linked to depression ^58^, (**Fig. 4E**) and *QPCT*, a pituitary glutaminyl-peptide cyclotransferase that has been previously associated with SZ ^59^ (**Fig. 4F**).

Genes that vary across donors were enriched for eQTLs detected in post mortem brain tissue ^48^ (**Fig. 4G)**, meaning that observed inter-individual expression variation reflected genetic regulation of expression. Conversely, genes with expression variation attributable to cell type (CTC scores) are either neutral or depleted for genes under genetic control, indicating that variation in CTC was either stochastic or epigenetic, but did not reflect genetic differences between individuals. Finally, the high percentage of residual variation not explained by factors considered here suggests that there are other uncharacterized sources of expression variation, including stochastic canalization effects or unexplained variation in CTC.

### Coexpression analysis identifies modules enriched for SZ and CTC

Genes with similar functions are known to share regulatory mechanisms and so are often coexpressed^60^. We used weighted gene coexpression network analysis (WGCNA)^61^ to identify modules of genes with shared expression patterns (**Fig. 5**, **SI Table 7**). Genes were clustered into modules of a minimum of 20 genes, and each module was labeled with a color (**SI Fig. 13**). Genes that did not form strong clusters were assigned to the grey module. Analysis was performed separately in hiPSC-NPCs and hiPSC-neurons; each module was evaluated for enrichment of genes for multiple biological processes. Many modules were highly enriched for genes that were significantly correlated with CTC scores at FDR < 5%, underscoring the genome-wide effects of cell type heterogeneity Genes that were differentially expressed between cases and controls in this study (see below) were enriched in the grey modules in both hiPSC-NPCs (OR=1.99, p<1.45e-5) and hiPSC-neurons (OR = 3.44, p< 5.04e-12, hypergeometric test), indicating that in this dataset, differentially expressed genes did not form a coherent structure but are instead widely distributed. While genes identified by genetic studies (i.e. common variants, CNVs, rare loss of function and de novo variants) showed moderate enrichment in many modules, they did not strongly overlap with the modules enriched for differentially expressed genes from this study; genes that were differentially expressed in the CommonMind Consortium post mortem dataset^48^ showed less of an enrichment signal. Finally, gene sets corresponding to the neural proteome show the strongest enrichment in the brown module from hiPSC-neurons, including, the targets of FMRP (OR = 4.06, p<2.84e-40) and genes involved in post-synaptic density (OR = 3.35, p<5.45e-22).

**Figure 5:**
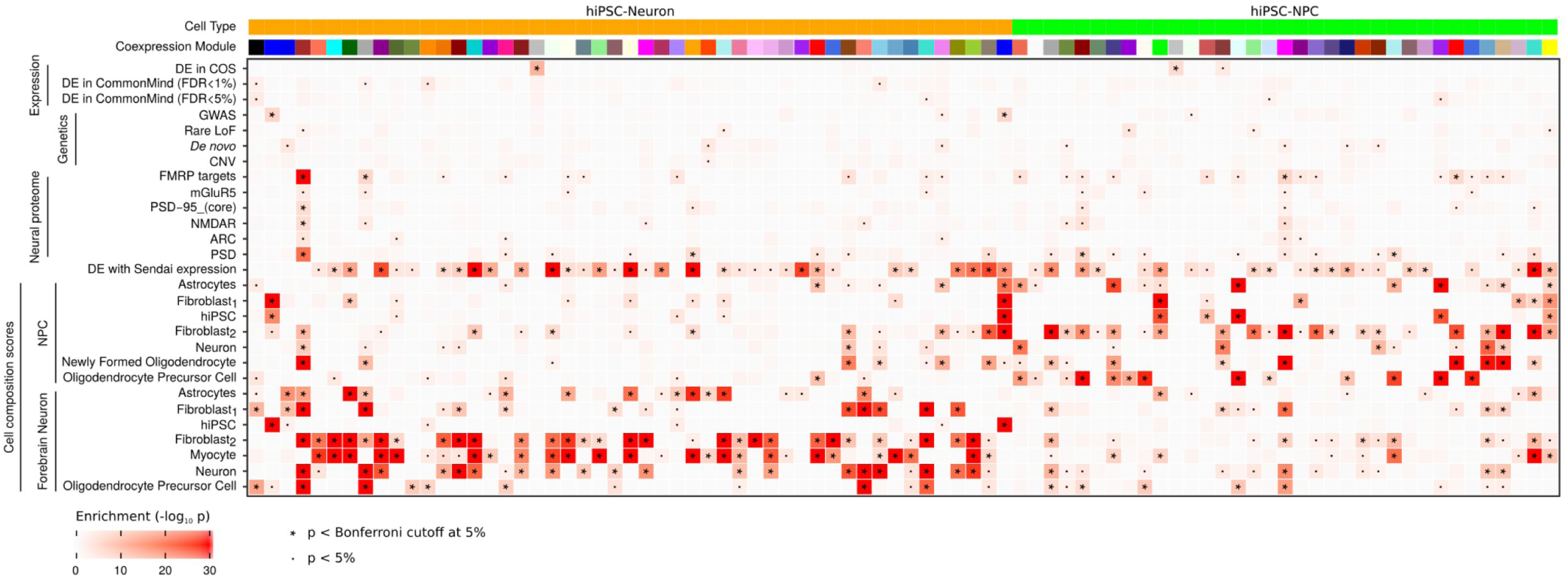
Clustering of genes into coexpression modules reveals module-specific enrichments. Enrichment significance (-log_10_ p-values from hypergeometric test) are shown for coexpression modules from hiPSC-NPCs and hiPSC-neurons. Each module is assigned a color and only modules with an enrichment passing the Bonferroni cutoff in at least one category is shown. Enrichments are shown for gene sets from RNA-Seq studies of differential expression between schizophrenia and controls; genetic studies of schizophrenia, neuronal proteome^48^; and cell composition scores from hiPSC-NPCs and hiPSC-neurons in this study. P-values passing the 5% Bonferroni cutoff are indicated by ‘*’, and p-values less than 0.05 are indicated with ‘.’.

### Differential expression between COS and control hiPSC-NPCs and hiPSC-neurons

The central objective of this study was to determine if a gene expression signature of SZ could be detected in an experimentally tractable cell culture model (**Fig. 6**). Due to the ‘repeated measures’ study design where individuals are represented by multiple independent hiPSC-NPC and -neuron lines, we used a linear mixed model by applying the duplicateCorrelation ^62^ function in our limma/voom analysis ^63,64^. This approach is widely used to control the false positive rate in studies of repeated measures ^65,66^ and its importance in hiPSC datasets was recently emphasized ^42^.

**Figure 6:**
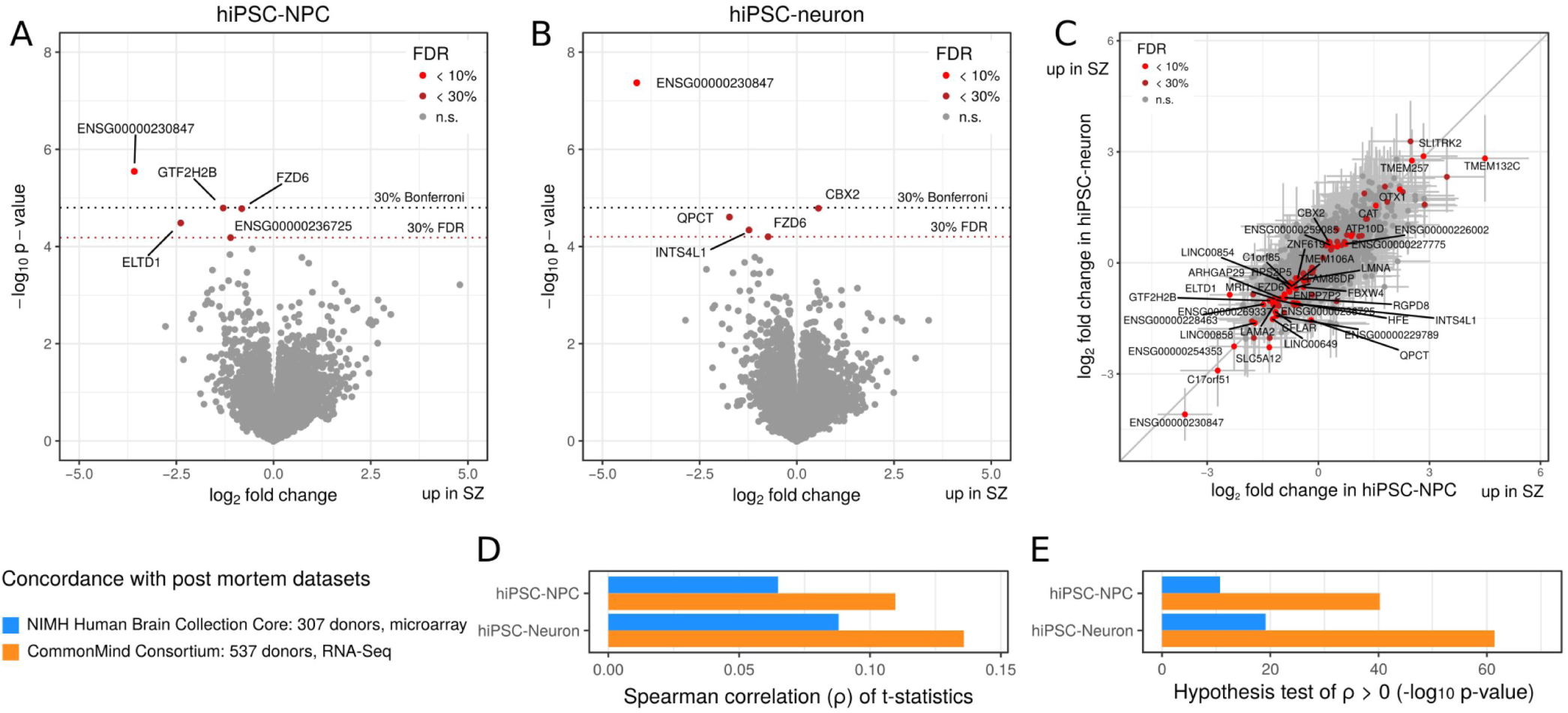
Differential expression between schizophrenia and controls. **A,B**) Volcano plot showing log_2_ fold change between cases and controls and the – log_10_ p-value for each gene in **A**) hiPSC-NPC and **B**) hiPSC-neuron samples. Genes are colored based on false discovery rate: light red (FDR < 10%), dark red (FDR < 30%), grey (n.s.: not significant). Names are shown for genes with FDR 30%. Dotted grey line indicates Bonferroni cutoff corresponding to a p-value of 0.30. Dashed dark red line indicates FDR cutoff of 30% computed by qvalue (Storey, 2002). **C**) Log_2_ fold change between cases and control in hiPSC-NPCs (x-axis) compared to log_2_ fold change between cases and controls in hiPSC-neurons (y-axis). Genes are colored according to differential expression results from combined analysis of both cell types: light red (FDR < 10%), dark red (FDR < 30%), grey (n.s.: not significant). Error bars represent 1 standard deviation around the log_2_ fold change estimates. **D,E**) Analysis of concordance between differential expression results of schizophrenia versus controls from the current study and two adult post mortem cohorts ^48^. Concordance is evaluated based on spearman correlation between t-statistics from two datasets. **D**) Spearman correlation between t-statistics from the current study (from hiPSC-NPCs and-neurons) and the two post mortem cohorts. **E**)-log_10_ p-values from a one-sided hypothesis test for the Spearman correlation coefficients from (**D**) being greater than zero.

Differential expression analysis between cases and controls in hiPSC-NPCs (**Fig. 6A**) identified 1 gene with FDR < 10% and 5 genes with FDR < 30%; analysis in hiPSC-neurons (**Fig. 6B**) identified 1 gene with FDR < 10% and 5 genes with FDR < 30% (**SI Table 8**).

While plausible candidates such as *FZD6* and *QPCT* were differentially expressed, gene set enrichment testing did not implicate a coherent set of pathways (**SI Table 9**). Since SZ is a highly polygenic disease ^67,68^ and this dataset is underpowered due to the small sample size ^48^, we expected the disease signal to be subtle and distributed across many genes. Despite performing extensive analysis using sophisticated statistical methods ^69-71^ built on top of the limma/voom framework ^63^ that incorporated genes that were not genome-wide significant and using permutations to empirically set the significance cutoff (see Methods), we failed to identify a coherent biological enrichment. Nonetheless, there was an unexpected concordance in the differential expression analysis between COS and control hiPSC-NPCs and hiPSC-neurons, which showed remarkably similar log_2_ fold changes (**Fig. 6C**). Moreover, no genes had log_2_ fold changes that were statistically different in the two cell types, although we were underpowered to detect such differences.

Overall, our differential expression analysis demonstrated that case-control hiPSC-based cohorts remain under-powered to resolve biologically coherent SZ-associated processes. Nonetheless, the concordance in the disease signature identified in hiPSC-NPCs and hiPSC-neurons implies that future studies could focus on just one cell type.

### Concordant differential gene expression in case-control hiPSC-NPCs and hiPSC-neurons with two much larger post mortem datasets

While it is well-understood that all hiPSC-based studies of SZ remain under-powered due to small sample sizes and polygenic disease architecture, what is less appreciated is that post mortem approaches are similarly constrained. Using allele frequencies from the Psychiatric Genetics Consortium data, the median number of subjects needed to obtain 80% power to resolve genome-wide expression differences in SZ cases was estimated to be ∼28,500, well beyond any existing data set ^48^. Nonetheless, we evaluated the concordance of our dataset with the findings of two much larger post mortem studies (CommonMind Consortium (CMC): RNA-Seq from 537 donors; NIMH Human Brain Collection core (HBCC), microarrays from 307 donors) by computing the correlation in t-statistics from the differential expression analysis between cases and controls.

The Spearman correlation between our hiPSC-NPC results and the CMC and HBCC results were 0.108 and 0.0661, respectively; for the hiPSC-neurons results, the correlations were 0.134 and 0.0896, respectively (**Fig. 6D**, **SI Fig. 14-15**). These correlations were highly statistically significant (**Fig. 6E**) for both hiPSC-NPCs: p < 4.6e-40 and 7.8e-12 for CMC and HBCC, respectively; and for hiPSC-neurons: p < 6.7e-61 and 1.6e-20 respectively (Spearman correlation test). Similar results were obtained by using Pearson correlation and by evaluating the concordance using the log_2_ fold changes from each dataset (**SI Fig. 14-15**). This stronger concordance of hiPSC-neurons (relative to hiPSC-NPCs) with post mortem findings is consistent with the hypothesis that neurons are the cell type most relevant to SZ risk ^72^, but our ability to resolve it is perhaps surprising in that neurons are estimated to comprise a minority of the cells in brain homogenate ^73^.

While the concordance with CMC was observed when correcting for any set of CTC scores (or none), the concordance with HBCC was only apparent when correcting for the fibroblast_1_ CTC score (**SI Fig. 16**). This illustrates the importance of accounting for CTC and the fact that concordance can be obscured by biological sources of expression variation. The genes for which the differential expression signal was boosted by accounting for the fibroblast_1_ score were enriched for brain and synaptic genesets, including specific biological functions such as FMRP and mGluR5 targets (**SI Fig. 17,18**).

This result indicates that although the concordance between hiPSC-NPCs and hiPSC-neurons with two post mortem datasets is relatively low due to the small sample size and low power of our current study, the concordance of the biological findings will increase with increasing sample size in future studies.

## DISCUSSION

SZ is a complex genetic disease arising through a combination of rare and common variants. Recent large-scale genotyping studies have begun to reveal the extent to which SZ risk reflects rare copy number variants (CNVs) ^30^ and coding mutations ^74^, as well as common single nucleotide polymorphisms (SNPs) with small effect sizes ^68^. The strongest finding to date from these genetic studies is that SZ-associated variants are enriched for pathways primarily associated with synaptic biology ^74,75^. Although more than 50 post mortem gene expression studies of SZ have been reported, the results have been inconsistent, likely owing to the small sample sizes involved ^48^. The largest of these, comparing brain tissue from 258 subjects with SZ and 279 controls did not find evidence for case–control differential expression among the implicated SZ risk genes; moreover, by modeling both the allele frequencies and the predicted allelic effects on gene expression, they predicted the median number of subjects needed to obtain genome-wide power (80%) to be ∼28,500 ^48^. This issue of small sample sizes is not unique to post mortem studies, and may be exacerbated in hiPSC-based experiments through the variability that arises as a result of the reprogramming and differentiation processes. We established an hiPSC cohort of COS patients ^76-80^, testing our ability to model gene expression changes associated with both common and rare variants *in vitro*. While other studies have focused on SZ cohorts comprised of relatively few individuals with rare mutations ^4,5,19^, we sought to determine to what extent a larger cohort captured the expression signature of polygenic SZ, focusing on COS due to the higher genetic burden of both rare and common variants in these patients.

The goal of studying patient-derived cell culture models is to develop an experimentally tractable platform that recapitulates a donor-specific gene expression signature. Retaining this donor-specific signature is essential to studying case control differences. In two recent studies of hiPSCs, variance across donors explained a median of ∼6% ^55^ and 48.8% ^35^ of expression variation, while the effect of donor was much smaller (2.2%) in this study. We hypothesize that donor effects are reduced due to stochastic noise in the differentiation from hiPSCs to neurons; it remains to be established whether different hiPSC-derived cell types will retain more or less donor signal over the course of differentiation. In our dataset, while genes with high expression variation across donors were enriched for eQTLs detected in post mortem brain, substantial expression variation within donors obscured some biological signal. In order to identify biological or technical variations that explained this intra-donor expression variation, we implemented a quality control pipeline to detect sample mislabeling, cell culture contamination, residual Sendai virus expression, incomplete X-inactivation and batch effects in sample processing; however, it was only accounting for variation in CTC that significantly decreased intra-donor variation.

Given the challenges of low statistical power, substantial intra-donor variation, and the range of complicating factors that can obscure the disease signal, future hiPSC-based studies of human disease should be carefully designed to maximize power. One particular challenge affecting many studies is the tradeoff between increasing the number of biological replicates and increasing the number of donors. The statistical concept of ‘effective sample size’ (ESS) addresses this issue directly and indicates that the tradeoff is dependent on the cost per donor and per hiPSC line in addition to the fraction of expression variation explained by donor (**Supplementary Text**). When a study includes multiple correlated samples from the same donor, the ESS is defined as the sample size of a study with equivalent power composed of only independent samples (**Fig. 7**). When the cost for each donor and each additional replicate are equal, adding an additional donor will increase the ESS by one unit (**Fig. 7A**), while adding an additional sample from an existing donor will increase the ESS by only a fraction of a unit because a sample correlated with it is already in the dataset. The contribution of each addition sample is determined by the donor effect. Therefore, when biological replicates from the same donor are very correlated, the increase in ESS can be small. Conversely, adding replicates when there is high intra-donor variability (i.e. a low donor effect) can have a larger increase on ESS. The fact that the donor effect in the current study is lower than in previous hiPSC studies^35,55^ affects the contribution of each additional sample to the ESS (**Fig. 7B**). When the costs for an additional hiPSC line are less than the cost of an additional donor, the calculus changes in favor of including additional biological replicates (**Fig. 7C,D**). We have developed a public website (gabrielhoffman.shinyapps.io/design_ips_study/) that computes the ESS in order to design a study to maximize power. These calculations consider constraints on either total budget or number of donors, as the relative cost and donor effect change. Overall, our conclusion is that the best way to maximize ESS, while controlling the false positive rate, is often to use one hiPSC line per donor and increase the number of donors, rather than using multiple replicate clones from a smaller set of donors ^42,65,66^.

**Figure 7:**
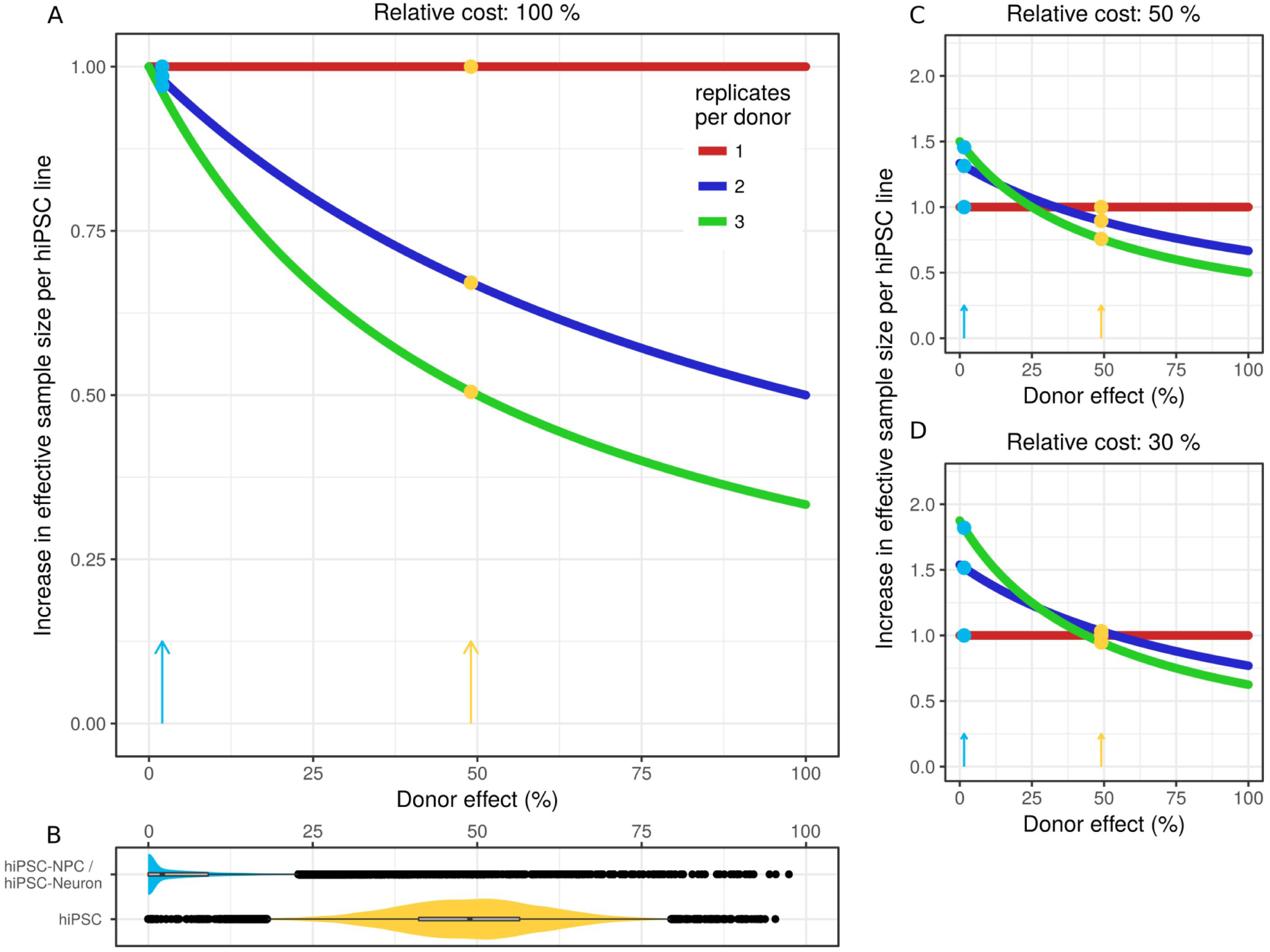
Maximizing power in hiPSC studies depends on relative costs and the fraction of expression variation across donors. **A)** The increase in effective sample size (ESS) for each additional hiPSC line added to the dataset shows as a function of the donor effect when the cost or an additional hiPSC line is the same as the cost for an additional donor. The increase in ESS is constant for the first replicate from a donor, while the contribution of the second or third replicates depend heavily on the donor effect. Colored points and arrows indicate the increase in ESS based on the donor effect from the current study (blue) and hiPSCs (orange)^35^. **B)** Violin plots show the full distribution and median donor effect computed by variancePartition for the current study (blue) and hiPSCs (orange). The median values across all genes correspond to the colored arrows and points in the other panels. **C)** Plot of ESS as in (**A**) but where the relative cost of an additional hiPSC line is 50% of the cost of an additional donor. **D)** Plot of ESS as in (**A**) but where the relative cost of an additional hiPSC line is 30% of the cost of an additional donor.

In addition to maximizing cohort ESS, future studies will benefit from decreasing intra-donor expression variation by optimizing neuronal differentiation/induction protocols to focus on decreasing cellular heterogeneity (rather than increasing total yield). The generation of single cell sequencing datasets from hiPSC-NPCs and/or hiPSC-neurons will further yield a custom reference panel with which to improve CTC deconvolution. In fact, our results suggest that to maximize ESS while minimizing associated costs, it may be sufficient to focus on hiPSC-NPCs rather than hiPSC-neurons. Given our improved understanding of the challenges associated with studying highly polygenic diseases as well as the biological constraints encountered here, disease signal will be further improved by reducing disease heterogeneity through focusing on cohorts of patients with shared genetic variants and/or the genetic engineering of isogenic hiPSC lines to introduce or repair SZ-relevant variants.

Despite our relatively small sample size, we were able to identify a subtle but statistically significant concordance between both COS hiPSC-NPCs and hiPSC-neurons with two recent SZ post mortem cohorts ^48^, an effect that was strongest in hiPSC-neurons. This indicated that shared biological pathways were disrupted in our hiPSC dataset and the adult post mortem donors. Moving forward, increasing the sample size of hiPSC-based cohorts will only improve the concordance. The surest strategy to improve the power of case-control comparisons is to integrate a growing number of post mortem and hiPSC studies. To facilitate improved sharing between stem cell laboratories, all hiPSCs have already been deposited at the Rutgers University Cell and DNA Repository (study 160; www.nimhstemcells.org/). We urge widespread sharing of all RNA-Seq data and reproducible scripts, and so make ours available at www.synapse.org/hiPSC_COS.

## SI FIGURE LEGENDS

SI Figure 1: **Ribosomal RNA rate computed from each RNA-Seq experiment**

SI Figure 2: **Effect of copy number variation on expression of proximal genes.** Expression of genes near CNV breakpoints were plotted and z-score of expression of each gene was used to identify expression outliers. Each line presents the expression of the set of genes for individuals with the CNV (red) and without the CNV (grey). Z-scores are plotted at the midpoint of the body of each gene.

SI Figure 3: **Quality control for sex, contamination and mislabeling. A**) Check that labeled sex is concordance with gene expression on chrX, and chrY. Plot of the sum of expression of 6 chrY genes (*USP9Y*, *UTY*, *NLGN4Y*, *ZFY*, *RPS4Y1*, *TXLNG2P*) versus expression on *XIST* from chrX. Males (blue) have distinct expression patterns of high chrY and low chrX expression. High quality female samples (red) have high chrX expression and low chrY expression. Problematic samples (grey) have intermediate expression patterns due to problems in X-inactivation, sample mislabeling or contamination involving a male and female sample. These samples were excluded from further analysis. These individuals are not known to have Klinefelter’s or other sex chromosome abnormality that would produce this observation. **B**) Contamination analysis using VerifyBamID ^44^ comparing variants called for each sample from RNA-Seq to variants from PsychChip and whole exome sequencing of the donors. Individual 499 shows a contamination percentage of 100%, recapitulating a known issue with sample mislabeling. Sample 1275-B-3F has a contamination percentage of 50%, consistent with (A) where this sample shows and expression patter intermediate between male and female. This sample is likely contains both male and female RNA.

SI Figure 4: **Quantifying residual Sendai virus from RNA-Seq reads. A**) Analysis workflow illustrating *de novo* assembly with Trinity/Inchworm, aligning contigs to Sendai genome and quantifying Sendai expression for each RNA-Seq experiment. **B**) Plot from NCBI showing results of BLAST alignment to the Sendai virus genome of all *de novo* contigs compiled across all 94 RNA-Seq experiments. Notice that Sendai gene F is not observed in the dataset likely due to the fact that the virus used in the experimental procedure was engineered. **C**) Quantification of Sendai expression in counts per million for each RNA-Seq experiment.

SI Figure 5: **Genes differentially expressed based on residual Sendai virus expression. A**) Gene set enrichment based on hypergeometric test for genes with FDR < 5%. **B**) Differential expression results for 3 Yamanaka factors genes used in a Sendai virus vector in the hiPSC reprogramming. *POU5F1* (i.e. *OCT4*) is not expressed at sufficient levels to be included in this analysis.

SI Figure 6: **Comparing expression patterns in hiPSC-NPC and hiPSC-neurons. A**) Venn diagram indicating high overlap of genes expressed at log_2_ RPKM of 1 in each cell type. **B**) Jaccard similarity between sets of genes that are expressed in each cell type at a level exceeding the expression cutoff on the x-axis. This indicates high overlap between sets of expressed genes. **C**) Volcano plot showing -log_10_ p-value and log_2_ fold change between hiPSC-NPC and hiPSC-neurons. Genes with FDR < 1% are indicated in light red and genes with FDR < 5% are indicated in dark red. Remaining genes are show in grey. **D,E**) Gene set enrichment tests based on hypergeometric test for gene sets in MSigDB for genes with FDR < 1% in D) hiPSC-NPCs and E) hiPSC-neurons.

SI Figure 7: **Genes with high inter-donor expression variation in hiPSC-NPCs and -neurons are enriched for brain cis-eQTLs.** Fold enrichment (log_2_) for the 2000 top cis-eQTLs discovered in post mortem dorsolateral prefrontal cortex data generated by the CommonMind Consortium^48^ shown for the inter-donor variance component in hiPSC-NPCs and ‒neurons. Each line indicates the fold enrichment for genes with the fraction of variance explained exceeding the cutoff indicated on the x-axis. Shaded regions indicate the 90% confidence interval based on 10,000 permutations of the variance fractions. Enrichments are shown on the x-axis until less that 100 genes pass the cutoff.

SI Figure 8: **Similarity between RNA-Seq samples from the same donor within each cell type. A**) Hierarchical clustering of RNA-Seq samples before correcting for the two fibroblast cell type composition scores. **B,C**) Correlation between samples from different donors compared to the correlation between samples from the sample donor. P-value indicates one-sided Wilcoxon test. B) Correlations for hiPSC-NPCs before correction. C) Correlations for hiPSC-neurons before correction.

SI Figure 9: **Cell type composition scores for current study and hiPSC-NPC and hiPSC-neuron samples from external datasets.**

SI Figure 10: **Accounting for fibroblast cell type composition scores increases similarity between RNA-Seq samples from the same donor within each cell type. A,B**) Correlation between samples from different donors compared to the correlation between samples from the sample donor for A) hiPSC-NPCs and B) hiPSC-neurons. P-value indicates one-sided Wilcoxon test.

SI Figure 11: **Violin plots of the percentage of variance explained by each variable over all the genes for multiple biological and technical sources of variation.**

SI Figure 12: **No differences in cell type composition scores between cases and controls. A**) Cell type composition scores stratified by case/control status for hiPSC-neurons and hiPSC-NPCs. **B**) -log_10_ p-values for hypothesis test (two-sided Wilcoxon) for each boxplot in (A). Dotted line indicates p-value of 0.05 and dashed line indicates Bonferroni cutoff at 5%. No tests are significant at even the nominal cutoff.

SI Figure 13: **Coexpression analysis. A**) Metric of scale free network topology for hiPSC-NPC and hiPSC-neuron networks. Dashed line indicates the software threashold of 9 used in the analysis. **B,C**) Dendrogram and module assignments from expression analysis for B) hiPSC-neurons and C) hiPSC-NPCs.

SI Figure 14: **Concordance between case/control differential expression results from hiPSC-NPCs from the current study and two adult post mortem cohorts. A,B**) Concordance between t-statistics from hiPSC-NPCs and A) CommonMind and B) HBCC cohorts. **C,D**) Concordance between log_2_ fold change estimates from hiPSC-NPCs and A) CommonMind and B) HBCC cohorts. Dashed grey line indicates a slope of 1. Dark red line indicates best fit line based on observed data. Correlation between two datasets are summarized in terms of Pearson correlation (R) and Spearman correlation (rho), each with corresponding p-values.

SI Figure 15: **Concordance between case/control differential expression results from hiPSC-neurons from the current study and two adult post mortem cohorts. A,B**) Concordance between t-statistics from hiPSC-neurons and A) CommonMind and B) HBCC cohorts. **C,D**) Concordance between log_2_ fold change estimates from hiPSC-neurons and A) CommonMind and B) HBCC cohorts. Dashed grey line indicates a slope of 1. Dark red line indicates best fit line based on observed data. Correlation between two dataset are summarized in terms of Pearson correlation (R) and Spearman correlation (rho), each with corresponding p-values.

SI Figure 16: **Concordance of case/control differential expression signatures between current study and post mortem cohorts depends on correction for cell type composition scores. A,B**) Spearman correlation between t-statistics for case/control differential expression analysis from the current study compared to A) CommonMind and B) HBCC cohorts were cell type composition scores were included as a covariate in the regression model. NULL indicates a model with no score included. Note the large effect of including the fibroblast_1_ score in the concordance with the HBCC cohort. **C,D**) One-sided hypothesis test for the correlation analysis in the previous panels for C) CommonMind and D) HBCC cohorts.

SI Figure 17: **Correcting for fibroblast_1_ cell type composition score in test of case/control differential expression affects specific genes in hiPSC-NPCs. A**) Comparison of absolute value of t-statistics from differential expression analysis including the fibroblast_1_ score as a covariate compared to absolute t-statistics omitting it. Dashed line indicates a slope of 1. Genes are colored based on their difference between the two analyses. Red indicates the 500 genes with the greatest increase in the absolute t-statistic and blue indicates the 500 genes with the greatest decrease. The remaining genes are in black. **B**) Histogram of differences in absolute t-statistics from (A). Dashed lines indicate the cutoff for the 500 genes with greatest increase (red) and greatest decrease (blue). **C,D)** Gene set enrichments using a hyper geometric test for the 500 genes with the greatest C) increase and D) decrease of absolute t-statistics.

SI Figure 18: **Correcting for fibroblast_1_ cell type composition score in test of case/control differential expression affects specific genes in hiPSC-neurons. A**) Comparison of absolute value of t-statistics from differential expression analysis including the fibroblast_1_ score as a covariate compared to absolute t-statistics omitting it. Dashed line indicates a slope of 1. Genes are colored based on their difference between the two analyses. Red indicates the 500 genes with the greatest increase in the absolute t-statistic and blue indicates the 500 genes with the greatest decrease. The remaining genes are in black. **B**) Histogram of differences in absolute t-statistics from (A). Dashed lines indicate the cutoff for the 500 genes with greatest increase (red) and greatest decrease (blue). **C,D)** Gene set enrichments using a hyper geometric test for the 500 genes with the greatest C) increase and D) decrease of absolute t-statistics.

## SI TABLE LEGENDS

SI Table 1: **Clinical and laboratory information about each individual and sample**

SI Table 2: **Clinical and laboratory metadata used bioinformatics analysis**

SI Table 3: **Quality control statistics for RNA-Seq data**

SI Table 4: **Biotype counts for expressed genes**

SI Table 5: **Differential expression analysis based on residual Sendai virus expression**

SI Table 6: **Gene set enrichments for residual Sendai virus differential expression analysis**

SI Table 7: **Coexpression modules and gene set enrichments**

SI Table 8: **Differential expression analysis between SZ and controls**

SI Table 9: **Gene set enrichments for cell type composition differential expression analysis**

## AUTHOR CONTRIBUTIONS

K.J.B., B.J.H, G.E.H., P.S. contributed to experimental design. K.J.B, B.J.H, I.L completed all cell culture experiments. E.F. conducted microscopy experiments. P.G and J.R. developed the cohort. D.R. and E.A.S. analyzed genetic data. G.E.H. performed RNA-Seq analysis. K.J.B. and G.E.H. wrote the manuscript.

## ACKNOWLEDGMENTS

Kristen J Brennand is a New York Stem Cell Foundation - Robertson Investigator. The Brennand Laboratory is supported by a Brain and Behavior Young Investigator Grant, National Institute of Health (NIH) grants R01 MH101454 and R01 MH106056 and the New York Stem Cell Foundation. The Sklar Laboratory is supported by NIH grant R01 MH109897. We thank the FACS core at Icahn School of Medicine at Mount Sinai. This work was supported in part through the computational resources and staff expertise provided by Scientific Computing at the Icahn School of Medicine at Mount Sinai.

Thanks to Gang Fang, Laura Huckins, Noam Beckmann and David Panchision for critical reading of the manuscript.

## ABBREVIATIONS

schizophrenia, SZ; childhood onset schizophrenia, COS; human induced pluripotent stem cell, hiPSC; neural progenitor cell, NPC; genome wide association study, GWAS; copy number variation, CNV; single nucleotide polymorphism, SNP; expression quantitative trait loci, eQTL.

## REFERENCES

1. Marchetto, M.C. et al. A model for neural development and treatment of rett syndrome using human induced pluripotent stem cells. Cell 143, 527-39 (2010).

2. Pasca, S.P. et al. Using iPSC-derived neurons to uncover cellular phenotypes associated with Timothy syndrome. Nat Med 17, 1657-62 (2011).

3. Shcheglovitov, A. et al. SHANK3 and IGF1 restore synaptic deficits in neurons from 22q13 deletion syndrome patients. Nature 503, 267-71 (2013).

4. Lin, M. et al. Integrative transcriptome network analysis of iPSC-derived neurons from schizophrenia and schizoaffective disorder patients with 22q11.2 deletion. BMC Syst Biol 10, 105 (2016).

5. Yoon, K.J. et al. Modeling a genetic risk for schizophrenia in iPSCs and mice reveals neural stem cell deficits associated with adherens junctions and polarity. Cell Stem Cell 15, 79-91 (2014).

6. Marchetto, M.C. et al. Altered proliferation and networks in neural cells derived from idiopathic autistic individuals. Mol Psychiatry 22, 820-835 (2017).

7. Mariani, J. et al. FOXG1-Dependent Dysregulation of GABA/Glutamate Neuron Differentiation in Autism Spectrum Disorders. Cell 162, 375-90 (2015).

8. Mertens, J. et al. Differential responses to lithium in hyperexcitable neurons from patients with bipolar disorder. Nature 527, 95-9 (2015).

9. Bavamian, S. et al. Dysregulation of miR-34a links neuronal development to genetic risk factors for bipolar disorder. Mol Psychiatry 20, 573-84 (2015).

10. Topol, A. et al. Dysregulation of miRNA-9 in a Subset of Schizophrenia Patient-Derived Neural Progenitor Cells. Cell Rep 15, 1024-36 (2016).

11. Brennand, K.J. et al. Modelling schizophrenia using human induced pluripotent stem cells. Nature 473, 221-5 (2011).

12. Topol, A. et al. Altered WNT Signaling in Human Induced Pluripotent Stem Cell Neural Progenitor Cells Derived from Four Schizophrenia Patients. Biol Psychiatry 78, e29-34 (2015).

13. Brennand, K. et al. Phenotypic differences in hiPSC NPCs derived from patients with schizophrenia. Mol Psychiatry 20, 361-8 (2015).

14. Srikanth, P. et al. Genomic DISC1 Disruption in hiPSCs Alters Wnt Signaling and Neural Cell Fate. Cell Rep 12, 1414-29 (2015).

15. Zhao, D. et al. MicroRNA Profiling of Neurons Generated Using Induced Pluripotent Stem Cells Derived from Patients with Schizophrenia and Schizoaffective Disorder, and 22q11.2 Del. PLoS One 10, e0132387 (2015).

16. Lee, I.S. et al. Characterization of molecular and cellular phenotypes associated with a heterozygous CNTNAP2 deletion using patient-derived hiPSC neural cells. NPJ Schizophrenia 1, 15019 (2015).

17. Yu, D.X. et al. Modeling hippocampal neurogenesis using human pluripotent stem cells. Stem Cell Reports 2, 295-310 (2014).

18. Robicsek, O. et al. Abnormal neuronal differentiation and mitochondrial dysfunction in hair follicle-derived induced pluripotent stem cells of schizophrenia patients. Mol Psychiatry 18, 1067-76 (2013).

19. Wen, Z. et al. Synaptic dysregulation in a human iPS cell model of mental disorders. Nature 515, 414-8 (2014).

20. Haggarty, S.J., Silva, M.C., Cross, A., Brandon, N.J. & Perlis, R.H. Advancing drug discovery for neuropsychiatric disorders using patient-specific stem cell models. Mol Cell Neurosci 73, 104-15 (2016).

21. Mariani, J. et al. Modeling human cortical development in vitro using induced pluripotent stem cells. Proc Natl Acad Sci U S A 109, 12770-5 (2012).

22. Pasca, A.M. et al. Functional cortical neurons and astrocytes from human pluripotent stem cells in 3D culture. Nat Methods 12, 671-8 (2015).

23. Qian, X. et al. Brain-Region-Specific Organoids Using Mini-bioreactors for Modeling ZIKV Exposure. Cell 165, 1238-54 (2016).

24. Nicholas, C.R. et al. Functional maturation of hPSC-derived forebrain interneurons requires an extended timeline and mimics human neural development. Cell Stem Cell 12, 573-86 (2013).

25. Gordon, C.T. et al. Childhood-onset schizophrenia: an NIMH study in progress. Schizophr Bull 20, 697-712 (1994).

26. Rapoport, J.L., Giedd, J.N. & Gogtay, N. Neurodevelopmental model of schizophrenia: update 2012. Mol Psychiatry 17, 1228-38 (2012).

27. Rapoport, J.L., Addington, A.M., Frangou, S. & Psych, M.R. The neurodevelopmental model of schizophrenia: update 2005. Mol Psychiatry 10, 434-49 (2005).

28. Ahn, K. et al. High rate of disease-related copy number variations in childhood onset schizophrenia. Mol Psychiatry 19, 568-72 (2014).

29. Ahn, K., An, S.S., Shugart, Y.Y. & Rapoport, J.L. Common polygenic variation and risk for childhood-onset schizophrenia. Mol Psychiatry (2014).

30. Marshall, C. et al. A contribution of novel CNVs to schizophrenia from a genome-wide study of 41,321 subjects. bioRxiv (2016).

31. Xu, J. et al. Inhibition of STEP61 ameliorates deficits in mouse and hiPSC-based schizophrenia models. Mol Psychiatry (2016).

32. Topol, A., Tran, N.N. & Brennand, K.J. A guide to generating and using hiPSC derived NPCs for the study of neurological diseases. J Vis Exp, e52495 (2015).

33. Hook, V. et al. Human iPSC neurons display activity-dependent neurotransmitter secretion: aberrant catecholamine levels in schizophrenia neurons. Stem Cell Reports 3, 531-8 (2014).

34. Tang, X. et al. Astroglial cells regulate the developmental timeline of human neurons differentiated from induced pluripotent stem cells. Stem Cell Res 11, 743-57 (2013).

35. Carcamo-Orive, I. et al. Analysis of Transcriptional Variability in a Large Human iPSC Library Reveals Genetic and Non-genetic Determinants of Heterogeneity. Cell Stem Cell (2016).

36. Ma, H. et al. Abnormalities in human pluripotent cells due to reprogramming mechanisms. Nature 511, 177-83 (2014).

37. Mekhoubad, S. et al. Erosion of dosage compensation impacts human iPSC disease modeling. Cell Stem Cell 10, 595-609 (2012).

38. Nazor, K.L. et al. Recurrent variations in DNA methylation in human pluripotent stem cells and their differentiated derivatives. Cell Stem Cell 10, 620-34 (2012).

39. Young, M.A. et al. Background mutations in parental cells account for most of the genetic heterogeneity of induced pluripotent stem cells. Cell Stem Cell 10, 570-82 (2012).

40. Ruiz, S. et al. Analysis of protein-coding mutations in hiPSCs and their possible role during somatic cell reprogramming. Nat Commun 4, 1382 (2013).

41. Hussein, S.M. et al. Copy number variation and selection during reprogramming to pluripotency. Nature 471, 58-62 (2011).

42. Germain, P.L. & Testa, G. Taming Human Genetic Variability: Transcriptomic Meta-Analysis Guides the Experimental Design and Interpretation of iPSC-Based Disease Modeling. Stem Cell Reports 8, 1784-1796 (2017).

43. Tomoda, K. et al. Derivation conditions impact X-inactivation status in female human induced pluripotent stem cells. Cell Stem Cell 11, 91-9 (2012).

44. Jun, G. et al. Detecting and estimating contamination of human DNA samples in sequencing and array-based genotype data. Am J Hum Genet 91, 839-48 (2012).

45. Grabherr, M.G. et al. Full-length transcriptome assembly from RNA-Seq data without a reference genome. Nat Biotechnol 29, 644-52 (2011).

46. Schlaeger, T.M. et al. A comparison of non-integrating reprogramming methods. Nat Biotechnol 33, 58-63 (2015).

47. Consortium, G.T. Human genomics. The Genotype-Tissue Expression (GTEx) pilot analysis: multitissue gene regulation in humans. Science 348, 648-60 (2015).

48. Fromer, M. et al. Gene expression elucidates functional impact of polygenic risk for schizophrenia. Nat Neurosci 19, 1442-1453 (2016).

49. Kang, H.J. et al. Spatio-temporal transcriptome of the human brain. Nature 478, 483-9 (2011).

50. Choi, J. et al. A comparison of genetically matched cell lines reveals the equivalence of human iPSCs and ESCs. Nat Biotechnol 33, 1173-81 (2015).

51. Newman, A.M. et al. Robust enumeration of cell subsets from tissue expression profiles. Nat Methods 12, 453-7 (2015).

52. Zhang, Y. et al. An RNA-sequencing transcriptome and splicing database of glia, neurons, and vascular cells of the cerebral cortex. J Neurosci 34, 11929-47 (2014).

53. Treutlein, B. et al. Dissecting direct reprogramming from fibroblast to neuron using single-cell RNA-seq. Nature 534, 391-5 (2016).

54. Hoffman, G.E. & Schadt, E.E. variancePartition: interpreting drivers of variation in complex gene expression studies. BMC Bioinformatics 17, 483 (2016).

55. Kilpinen, H. et al. Common genetic variation drives molecular heterogeneity in human iPSCs. Nature 546, 370-375 (2017).

56. McKean, D.M. et al. FAK induces expression of Prx1 to promote tenascin-C-dependent fibroblast migration. J Cell Biol 161, 393-402 (2003).

57. Ocana, O.H. et al. Metastatic colonization requires the repression of the epithelial-mesenchymal transition inducer Prrx1. Cancer Cell 22, 709-24 (2012).

58. Wilkinson, M.B. et al. A novel role of the WNT-dishevelled-GSK3beta signaling cascade in the mouse nucleus accumbens in a social defeat model of depression. J Neurosci 31, 9084-92 (2011).

59. Ripke, S. et al. Genome-wide association analysis identifies 13 new risk loci for schizophrenia. Nat Genet 45, 1150-9 (2013).

60. Zhang, B. & Horvath, S. A general framework for weighted gene co-expression network analysis. Stat Appl Genet Mol Biol 4, Article17 (2005).

61. Langfelder, P. & Horvath, S. WGCNA: an R package for weighted correlation network analysis. BMC Bioinformatics 9, 559 (2008).

62. Smyth, G.K., Michaud, J. & Scott, H.S. Use of within-array replicate spots for assessing differential expression in microarray experiments. Bioinformatics 21, 2067-75 (2005).

63. Ritchie, M.E. et al. limma powers differential expression analyses for RNA-sequencing and microarray studies. Nucleic Acids Res 43, e47 (2015).

64. Law, C.W., Chen, Y., Shi, W. & Smyth, G.K. voom: Precision weights unlock linear model analysis tools for RNA-seq read counts. Genome Biol 15, R29 (2014).

65. Jostins, L., Pickrell, J.K., MacArthur, D.G. & Barrett, J.C. Misuse of hierarchical linear models overstates the significance of a reported association between OXTR and prosociality. Proc Natl Acad Sci U S A 109, E1048 (2012).

66. Pinheiro, J. & Bates, D. Mixed-effects models in S and S-Plus, (Springer, New York, 2000).

67. Loh, P.R. et al. Contrasting genetic architectures of schizophrenia and other complex diseases using fast variance-components analysis. Nat Genet 47, 1385-92 (2015).

68. Schizophrenia Working Group of the Psychiatric Genomics Consortium. Biological insights from 108 schizophrenia-associated genetic loci. Nature 511, 421-7 (2014).

69. Majewski, I.J. et al. Opposing roles of polycomb repressive complexes in hematopoietic stem and progenitor cells. Blood 116, 731-9 (2010).

70. Wu, D. et al. ROAST: rotation gene set tests for complex microarray experiments. Bioinformatics 26, 2176-82 (2010).

71. Wu, D. & Smyth, G.K. Camera: a competitive gene set test accounting for inter-gene correlation. Nucleic Acids Res 40, e133 (2012).

72. Skene, N.G. et al. Genetic Identification Of Brain Cell Types Underlying Schizophrenia. bioRxiv (2017).

73. Sherwood, C.C. et al. Evolution of increased glia-neuron ratios in the human frontal cortex. Proc Natl Acad Sci U S A 103, 13606-11 (2006).

74. Fromer, M. et al. De novo mutations in schizophrenia implicate synaptic networks. Nature 506, 179-84 (2014).

75. Purcell, S.M. et al. A polygenic burden of rare disruptive mutations in schizophrenia. Nature 506, 185-90 (2014).

76. Sporn, A. et al. 22q11 deletion syndrome in childhood onset schizophrenia: an update. Mol Psychiatry 9, 225-6 (2004).

77. Shaw, P. et al. Childhood-onset schizophrenia: A double-blind, randomized clozapine-olanzapine comparison. Arch Gen Psychiatry 63, 721-30 (2006).

78. McCarthy, S.E. et al. Microduplications of 16p11.2 are associated with schizophrenia. Nat Genet 41, 1223-7 (2009).

79. Gogtay, N. et al. Dynamic mapping of human cortical development during childhood through early adulthood. Proc Natl Acad Sci U S A 101, 8174-9 (2004).

80. Eckstrand, K. et al. Sex chromosome anomalies in childhood onset schizophrenia: an update. Mol Psychiatry 13, 910-1 (2008).

